# OMIDIENT: Multiomics Integration for Cancer by Dirichlet Auto-Encoder Networks

**DOI:** 10.1101/2025.07.02.662608

**Authors:** Negar Safinianaini, Niko Välimäki, Roman Bresson, Alexandra Gorbonos, Kristiina Rajamäki, Lauri A. Aaltonen, Pekka Marttinen

## Abstract

To achieve a more comprehensive understanding of cancer, novel computational methods are required for the integrative analysis of data from different molecular layers, such as genomics, transcriptomics, and epigenomics. Here, we present a novel multi-omics integrative method that performs unsupervised representation learning, referred to as OMIDIENT: multiOMics Integration for cancer by DIrichlet auto-ENcoder neTworks. OMIDIENT provides a natural framework for modeling sparse and compositional latent representations by employing a deep generative model, where the latent space is distributed as the product of Dirichlet distributions. Applied to five different cancers, we demonstrate that OMIDIENT outperforms the top state-of-the-art unsupervised multi-omics integrative analysis approaches in clustering, classification, and reconstruction of missing data using mRNA expression data, DNA methylation data, and microRNA expression data. Furthermore, we provide interpretability analyses for OMIDIENT that not only support its improved performance, but also offer valuable insights into the underlying structure captured by the learned representations.

## 1 Introduction

Cancer is a complex disease in which various biological alterations give rise to characteristic patterns; discovering those structures is bound to have clinical relevance and can subsequently improve cancer treatment. Due to the heterogeneous nature of cancer, it is vital to identify specific patterns and clinical treatment options for different patients. Therefore, it is of utmost importance to provide a space where one can efficiently analyze the multimodal high-dimensional data. Integrating multiomics facilitates a comprehensive understanding of cellular diversity by combining data from different molecular layers, such as genomics, transcriptomics, and epigenomics. Each omics layer captures a different aspect of cellular function, and integration allows us to model the complex, interconnected nature of biology more effectively than any single omics type alone.

Multi-omics integration has been extensively studied in cancer research Huang (2019); Singh (2019); Poirion et al. (2018); Xie (2019); Chaudhary et al. (2018), with various methods proposed. In MOGONET Wang et al. (2021a), a graph-based supervised learning is proposed, while in MOCSS Chen et al. (2023), an unsupervised—learning without access to the subtyping labels—approach is adopted. Classical approaches have also been explored, such as NEMO Rappoport et al. (2019), which employs a neighborhood-based clustering strategy, and DeFusion Wang et al. (2021b), which formulates multiomics integration as a non-negative matrix factorization problem. Each of these methods addresses a specific downstream task, such as cancer subtyping or survival analysis. However, given the complexity of multiomics data, the typically limited number of tumor samples, and the exploratory nature of subtyping—which often involves discovering previously unknown subtypes—we are particularly motivated to adopt an unsupervised learning approach. Building on the growing field of *representation learning* Bengio et al. (2014), our goal is to learn deep, hierarchically abstract features that capture shared and modality-specific biological patterns, where more abstract concepts are higher in the hierarchy and deeper in the neural network. By embedding biological complexity into structured latent spaces, these compact and informative representations improve performance in downstream tasks such as subtyping and the prediction of survival and drug response.

To address these challenges, we propose a multi-omic integration method for unsupervised representation learning, referred to as OMIDIENT: MultiOMics Integration for cancer by DIrichlet auto-ENcoder neTworks. OMIDIENT is a novel probabilistic model for learning latent representations of tumor samples. It is inspired by two state-of-the-art methods for representation learning from different fields: MOCSS, a method for multi-omic data integration Chen et al. (2023), which captures omics-specific and omics-shared representations, and Dirichlet variational autoencoder Joo et al. (2019), which naturally finds distinct groups among the samples. We generalize the Dirichlet variational autoencoder by modeling the latent representations with a *product* of Dirichlet distributions, allowing the model to group samples along multiple dimensions, by using distinct parts of the latent representations, providing a natural framework for learning sparse and compositional structure in the data. Our method improves cancer subtyping and reconstruction of missing data. Dividing features into distinct groups allows the model to capture structured biological or functional relationships, such as variability in gene sets, pathways, or phenotypes. In this way, the internal variation remains interpretable and biologically meaningful, which we validate by analyzing the interpretability of our method.

Recent methods have shown strong performance in unsupervised learning tasks. We evaluated OMIDIENT on five public cancer datasets: breast, colon, colorectal, liver, and kidney cancer, learning representations using mRNA expression, DNA methylation, and microRNA expression data. Across these datasets, OMIDIENT outperformed existing approaches in cancer subtyping, evaluated through clustering and classification based on the learned representations, and achieved superior performance in missing omic data reconstruction. OMIDIENT is designed to flexibly integrate additional modalities if available. The code is publicly available at: https://github.com/negar7918/multiomics.

## 2 Results

### 2.1 Framework of OMIDIENT

We introduce OMIDIENT, shown in Fig. 1, an unsupervised multi-omics integration framework for tumor representation learning—these representations can be used for downstream tasks in cancer analysis, such as subtyping. Specifically, we introduce a novel probabilistic model, referred to as ProdDirVae and illustrated in part A in Fig. 1, to capture *S* distinct groups of tumor characteristics. Each omic dataset is modelled by ProdDirVae to capture omic-specific representations. For shared omic representations, we use contrastive learning similarly to MOCSS Chen et al. (2023), since contrastive learning is a state-ofthe-art powerful machine learning tool for capturing shared representations of multiple modalities You et al. (2022). We conduct comprehensive ablation studies to demonstrate the necessity of different components in OMIDIENT. A major advantage of ProdDirVae is its robustness toward high-dimensionality of multiomics, given limited tumor samples.

**Figure 1:**
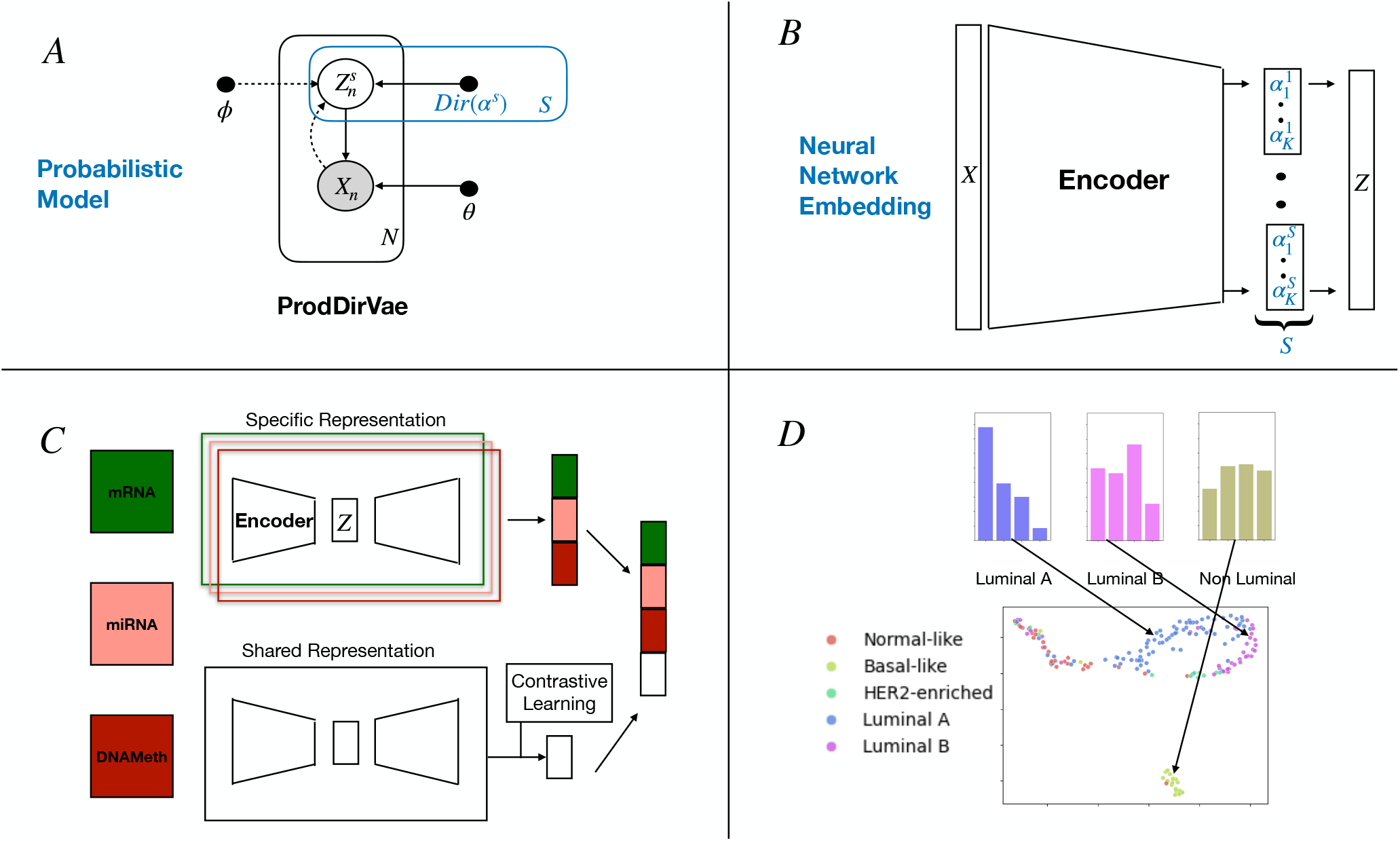
An overview of our proposed method. **A)** our novel probabilistic graphical model, which models the latent representation *Z* as belonging to *S* distinct groups. **B)** the corresponding unfolded computational graph of the neural network architecture. **C)** overview of learning omic-specific and shared representations. **D)** demonstration of the utility of the learned representation *Z*, where we perform BRCA subtyping based on *Z*. For clarity, we only show group *S* = 1 for mRNA omic, which clearly separates Luminal A and B subtypes, while the Basal-like subtype exhibits a near-uniform distribution, reflecting its non-luminal characteristics.

Motivated by the task of cancer subtyping, we apply clustering and classification using the learned tumor representations. In addition, we reconstruct each omic modality from these representations to assess their generative quality. For subtyping, we evaluate model performance using ground-truth labels across five public datasets: (1) breast invasive carcinoma (BRCA), (2) colon adenocarcinoma (COAD),(3) colorectal cancer (CRC), (4) liver hepatocellular carcinoma (LIHC), and (5) kidney renal clear cell carcinoma (KIRC). To demonstrate the effectiveness and robustness of OMIDIENT, we conduct extensive experiments across multiple clustering, classification, and missing data reconstruction tasks. Our analysis incorporates three omic types—mRNA expression, DNA methylation, and miRNA expression—capturing complementary and comprehensive views of tumor biology.

For the details of our method and the datasets, see Section Methods.

### 2.2 Clustering and Classification

For the downstream task of cancer subtyping, we compared the clustering performance of OMIDIENT with top unsupervised multi-omics integration frameworks, i.e., MOCSS Chen et al. (2023) a deep learning and contrastive learning framework, NEMO Rappoport et al. (2019) a neighborhood-based multi-omics clustering, DeFusion Wang et al. (2021b) a non-negative matrix factorization method, Concerto Yang et al. (2022) a contrastive learning tool, MultiVI Ashuach et al. (2023) a joint omic probabilistic model. Moreover, from machine learning literature, we adapted other probabilistic models similar to OMIDI-ENT to assess how variations in model design affect performance; namely, LassoAE Gao and et al (2025), VAE Kingma and Welling (2014), a variational autoencoder-based multi-omics integration, LapDirVae Srivastava and Sutton (2017), a Laplace approximated Dirichlet autoencoder-based multi-omics integration, and GamDirVae Joo et al. (2019), a Gamma approximated Dirichlet autoencoder-based multi-omics integration.

We evaluated the K-means clustering results using normalized mutual information (NMI), based on the ground-truth class labels provided for each dataset. We also applied the k nearest neighbors (k-NN) algorithm, reporting the classification accuracy in the latent space for methods that provided explicit representations Joo et al. (2019).

We evaluated all the methods on five datasets and six clustering and classification scenarios. BRCA data with five subtypes, COAD data with four subtypes, LIHC data with cancer and normal samples, KIRC with two subtypes, and CRC with four or two classes. Microsatellite instability (MSI) arises through defective DNA mismatch repair in the tumors, causing mutations particularly at repetitive DNA sequences called “microsatellites”; MSI classification system is relevant for CRC prognosis and treatment Taieb et al. (2022). Consequently, for the CRC with four classes, we used the following groups: 1) microsatellite instability (MSI), 2) microsatellite stability (MSS), 3) ambiguous tumors, and 4) normal samples. For the CRC with two classes, we used the following groups within consensus molecular subtype (CMS): 1) CMS1 group containing microsatellite instability immune, 14%, hypermutated, microsatellite unstable, and strong immune activation, and 2) other CMS groups and normal samples. For the details of the datasets and classes, see Section Methods.

Table 1 shows that our method outperforms all methods in unsupervised clustering of samples with the K-means algorithm on average (last column) and specifically on BRCA, KIRC, and CRC data. GamDirVae has the same performance as ours on KIRC data. The difference in performance on average is 5 per cent units higher than all other methods. As shown in Table 2, our method (last row) outperforms all methods also in terms of supervised classification of samples using the k-NN classifier. For each data, we achieve the highest or on par performance except for one grouping in the CRC dataset, which we study in more detail in the Section for Biological Interpretation of Learned Representations.

**Table 1:**
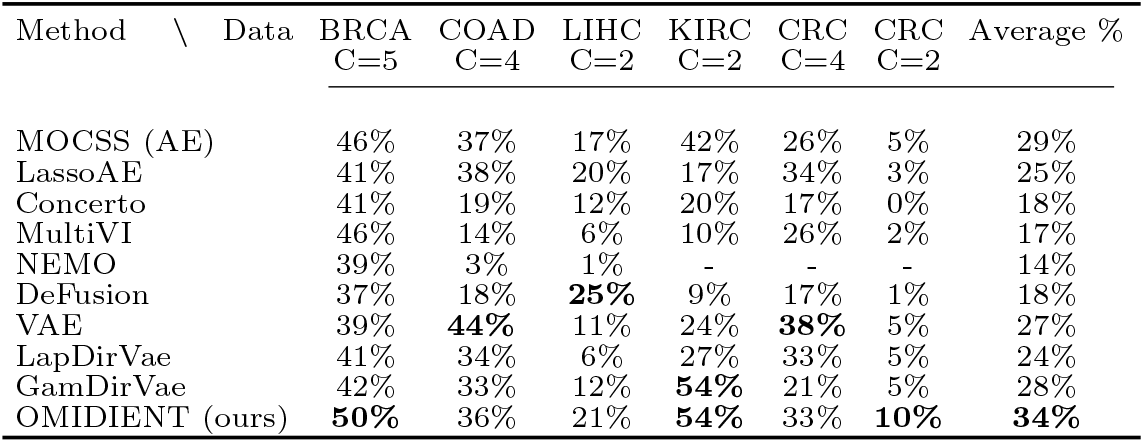
K-means performance, measured by NMI, for C classes based on the learned representations.

**Table 2:**
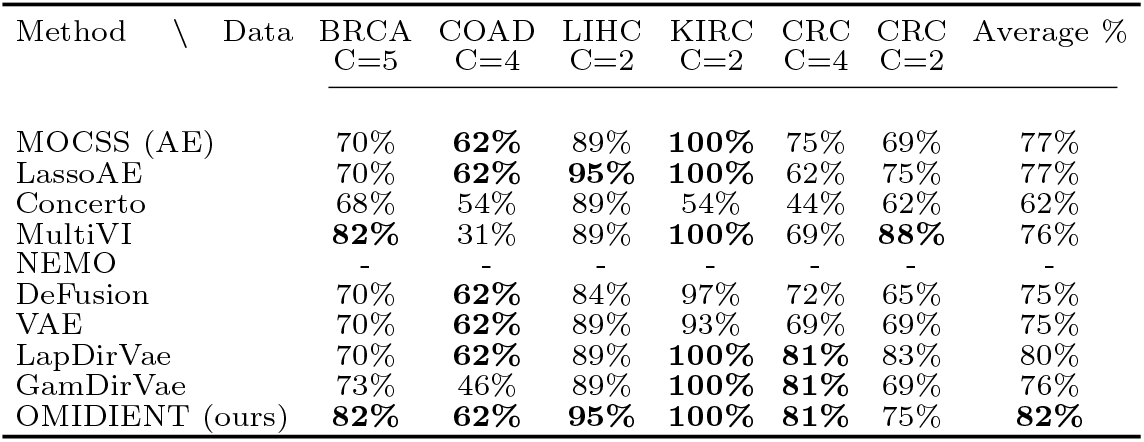
k-NN performance, measured by accuracy, for C classes based on the learned representations.

| Method \ Data | BRCA<br>C=5 | COAD<br>C=4 | LIHC<br>C=2 | KIRC<br>C=2 | CRC<br>C=4 | CRC<br>C=2 | Average % |
| --- | --- | --- | --- | --- | --- | --- | --- |
| MOCSS (AE) | 70% | **62%** | 89% | **100%** | 75% | 69% | 77% |
| LassoAE | 70% | **62%** | **95%** | **100%** | 62% | 75% | 77% |
| Concerto | 68% | 54% | 89% | 54% | 44% | 62% | 62% |
| MultiVI | **82%** | 31% | 89% | **100%** | 69% | **88%** | 76% |
| NEMO | - | - | - | - | - | - | - |
| DeFusion | 70% | **62%** | 84% | 97% | 72% | 65% | 75% |
| VAE | 70% | **62%** | 89% | 93% | 69% | 69% | 75% |
| LapDirVae | 70% | **62%** | 89% | **100%** | **81%** | 83% | 80% |
| GamDirVae | 73% | 46% | 89% | **100%** | **81%** | 69% | 76% |
| OMIDIANT (ours) | **82%** | **62%** | **95%** | **100%** | **81%** | 75% | **82%** |

Regarding NEMO, note that it does not provide any learned representations; hence, the k-NN classification task was not applicable. Moreover, NEMO could not be run on KIRC data due to its high dimensionality on the gene expression data. Finally, NEMO could not be executed on the CRC dataset due to an error in the original software, which we did not attempt to resolve within the scope of this study.

### 2.3 Reconstruction of Missing Data

We evaluated OMIDIENT on the task of missing data reconstruction, comparing it to MOCSS Chen et al. (2023), the top-performing state-of-the-art method as identified in Tables 1 and 2.

To simulate missing data scenarios, we conducted two experiments: in one, we completely omitted miRNA data from the test set; in the other, we randomly removed features across all three omics types. We used correlation to evaluate how close the reconstructed omics are to the original omics.

We evaluated our method and MOCSS on four datasets: BRCA data, COAD data, LIHC data, and KIRC. Table 3 shows that our method consistently outperforms MOCSS across all datasets and missing data scenarios. Fig. 2 presents the correlation plots comparing our method (OMIDIENT) and MOCSS, demonstrating that our approach achieves approximately 10 percentage points higher reconstruction accuracy.

**Table 3:** Reconstruction of missing miRNA omic and random features results.

| Method \ Data | BRCA |  | COAD |  | LIHC |  | KIRC |  |
| --- | --- | --- | --- | --- | --- | --- | --- | --- |
|  | miRNA | random | miRNA | random | miRNA | random | miRNA | random |
| MOCSS (AE) | 89% | 93% | 98% | 96% | 97% | 95% | 96% | 91% |
| OMIDIENT (ours) | **99%** | **99%** | **99%** | **99%** | **98%** | **100%** | **98%** | **100%** |

**Figure 2:**
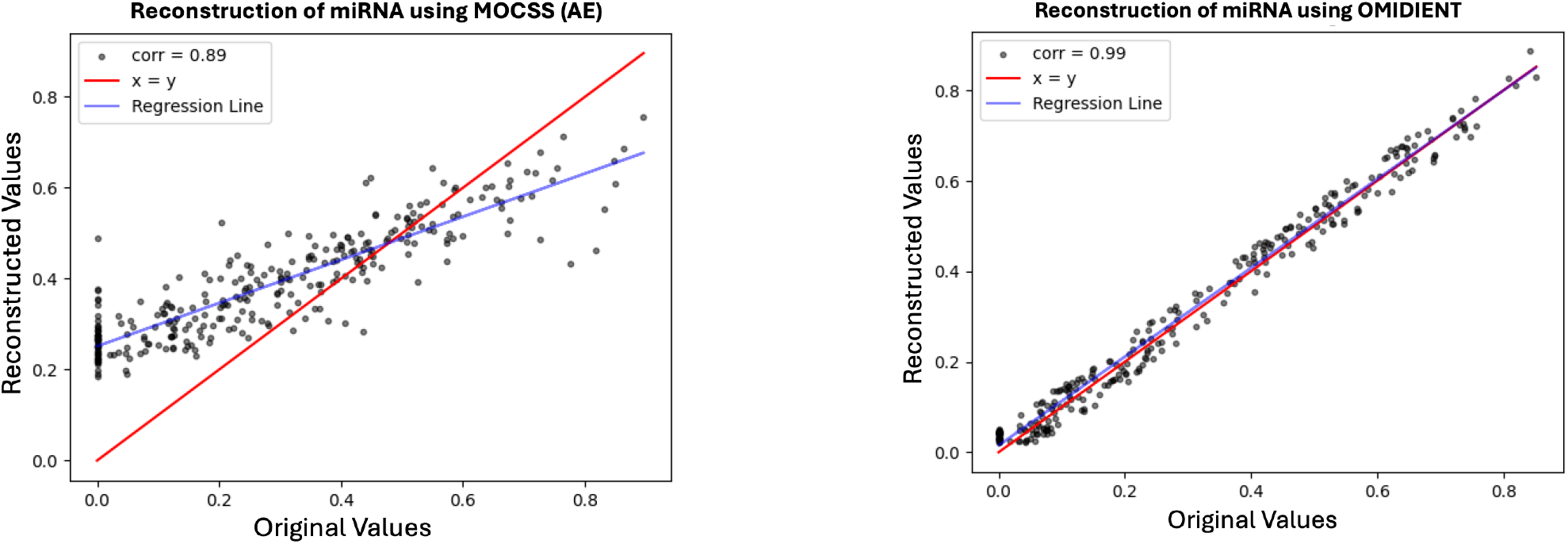
Reconstruction of BRCA data is compared between OMIDIENT (ours) and MOCSS on BRCA when miRNA omic is missing.

### 2.4 Interpretability

#### 2.4.1 Biological Interpretation of Learned Representations

In this section, we consider biological interpretations of the learned representations for the CRC data, since for those data, we have access to multiple annotations.

First, we interpret the learned representations in terms of the MSI and MSS grouping that are established subtypes in colorectal cancer Roepman (2014); Budinska (2013); Schlicker (2012); Sadanandam (2013); De Sousa E Melo (2013); Marisa (2013). In Fig. 3, we compare the UMAP illustration of the samples’ learned representations by the top models in Table 1 and 2, i.e., MOCCS (AE), VAE, LapDirVae, and OMIDIENT (ours). The samples include both tumors and normal colon samples, i.e., entirely nonneoplastic tissue. The axes x and y of each plot correspond to the two components of the UMAP transformation. Although our method (OMIDIENT) and LapDirVae show similar performance on k-means and k-NN metrics for the CRC data, the UMAP visualization reveals that tumor samples form denser and more distinctly separated clusters with our approach. While additional genomic or clinical annotations are needed to fully interpret these groups, the results suggest that our method may uncover novel subgroups, particularly within the MSS subtype.

**Figure 3:**
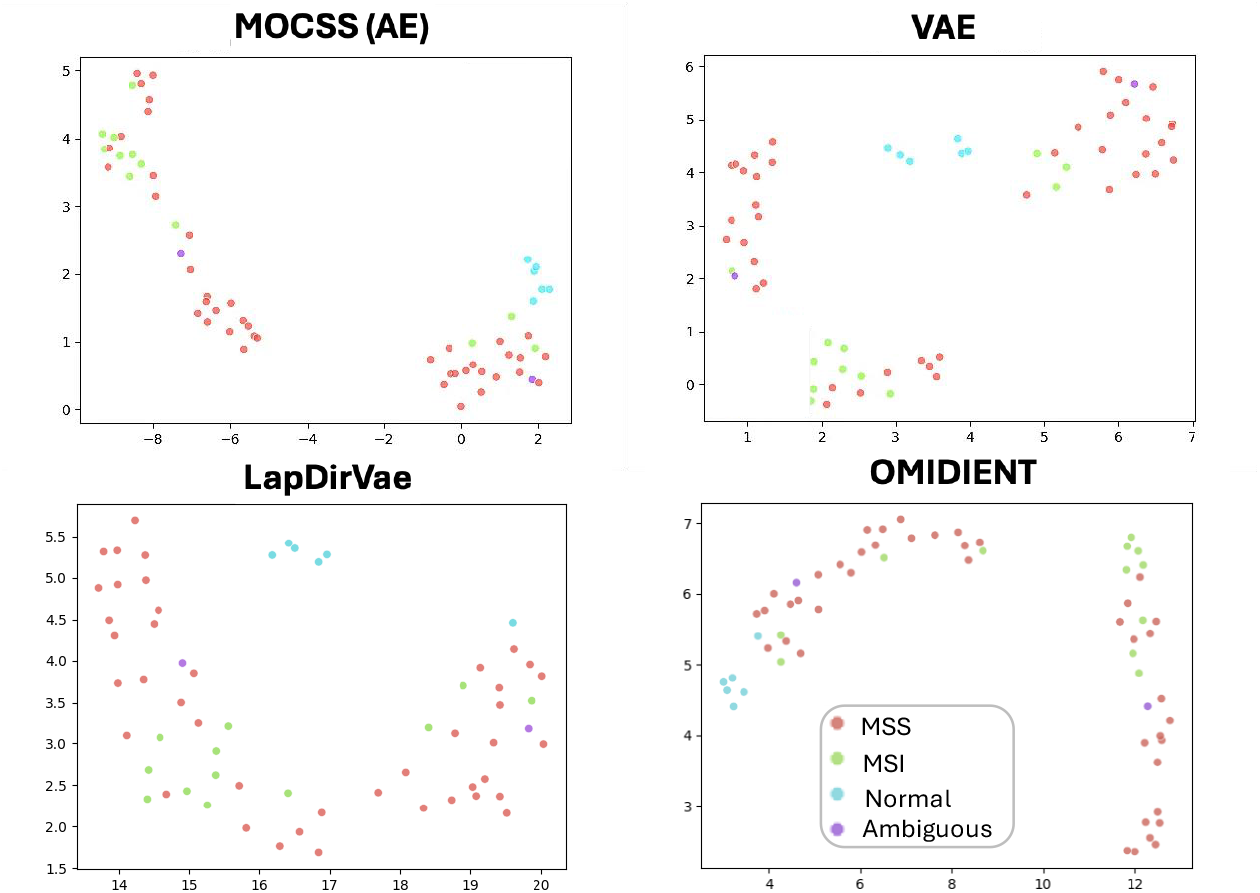
The samples’ learned representations by MOCSS (AE), VAE, LapDirVae, and Ours (OMIDI-ENT), which are mapped to UMAP, are shown.

Next, we interpret the learned representations by CMS groupings, a robust gene expression-based CRC classification system associated with biological mechanisms of tumorigenesis and with patient prognosis Guinney et al. (2015). Namely we compare CMS1 versus other CMS subtypes Guinney et al. (2015) because it most closely corresponds to the MSI subtype, i.e. most MSI tumors tend to be CMS1. In Fig. 4, we present UMAP visualizations of the learned representations from the top-performing models in Tables 1 and 2, namely MOCSS (AE), LapDirVae, MultiVI, and our method, OMIDIENT. Our method achieves the highest NMI score, OMIDIENT (bottom right), demonstrating a more distinct and well-separated grouping of CMS1 samples (highlighted in red) in the top-right region of the plot. In contrast, MOCSS (top left) places the CMS1 samples in close proximity to, or overlapping with, other CMS groups, indicating less discriminative representations. Finally, although MultiVI and LapDirVae achieve high accuracy in Table 2, the UMAP plots (bottom left and top right) reveal poor clustering of CMS1 (highlighted in red), suggesting that the seemingly high accuracy may be an artifact caused by class imbalance. This is further supported by its low NMI score.

**Figure 4:**
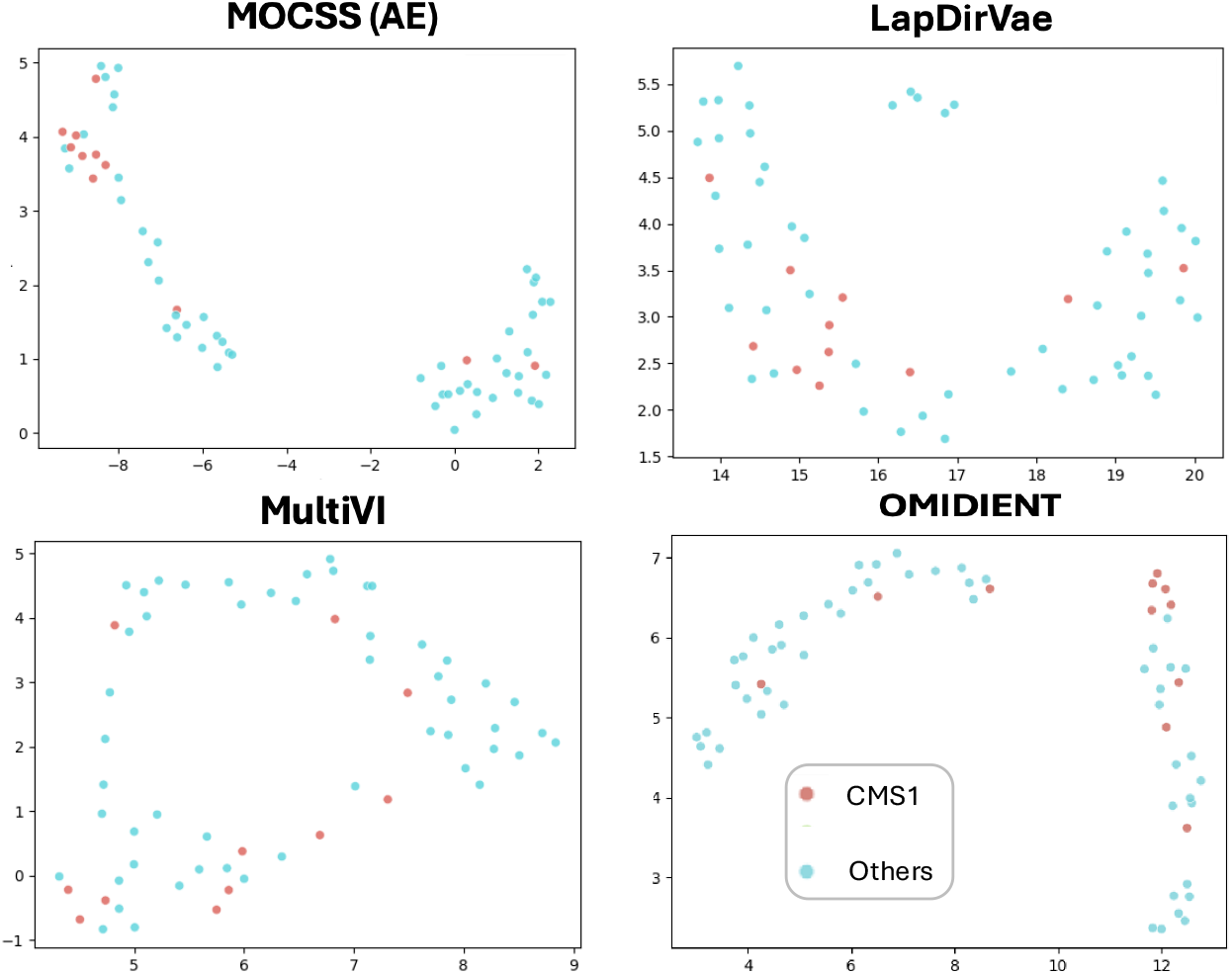
The samples’ learned representations by OMIDIENT, MOCSS (AE) and LapDirVae, which are mapped to UMAP are shown. CMS1 tumors, colored in red, are best clustered in OMIDIENT. This is also confirmed via NMI in Table 1.

Moreover, we annotate the samples by gender and tumor stages in Fig. 5. We can observe that our method distinguishes the genders clearly. Note that, due to the use of UMAP for dimensionality reduction, not all learned dimensions are visualized. As a result, some female samples appear on the left-hand side (LHS), representing normal, non-tumor tissue. Comparing this plot to Fig. 3, we can observe that “normal-adjacent” (male) tumor cluster in OMIDIENT has fewer MSI tumors and the “normal-distant” (female) tumor cluster has more MSI tumors. This observation is consistent with the well-established finding that MSI is more prevalent in colorectal cancers (CRCs) among females than males. It also appears that in Fig. 3, OMIDIENT can distinguish the MSI subgroup among women quite well into its own half of the female cluster, while in the male cluster MSI tumors are fully scattered around. In the figure on the right side, the tumor stages are annotated and the Adenomas are clustered in the middle of the right cluster, being close to stages I and II, which is sensible.

**Figure 5:**
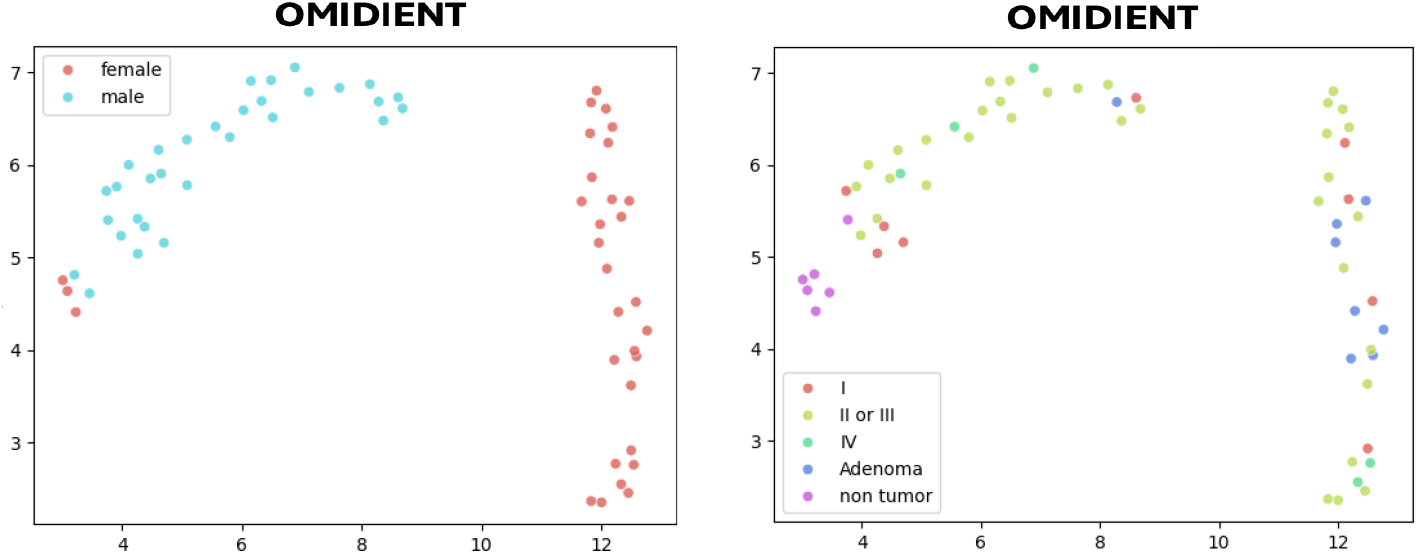
The samples’ learned representations by OMIDIENT. In the LHS, the gender is annotated. In the RHS, tumor stages are annotated, where non-tumor refers to normal samples and Adenoma to benign tumor samples.

In addition to the annotations above, we perform pathway enrichment analysis. Figure 6 shows the associations between latent representation values and different annotations. The shared representation associate with gender, tissue and TMN stage. The omic-specific representations make the most significant contributions to gene set enrichment analysis of key signaling pathways. The representations for mRNA and DNA methylation appear to be complementary and partially represent orthogonal signals from the two different omics. There were no associations with age at surgery. Some signaling pathways are known to be involved in different molecular subtypes of CRC Guinney et al. (2015), with JAK-STAT, VEGF and Wnt given here as a representative example. The gene set mRNA enrichment analysis showing CRC canonical pathways in tumor samples is inferred by using clusterProfiler (v4.10.0; org.Hs.eg.db v3.18.0), KEGG (accessed on May 15, 2025), and genes ranked by their gene expression correlation with each latent representation.

**Figure 6:**
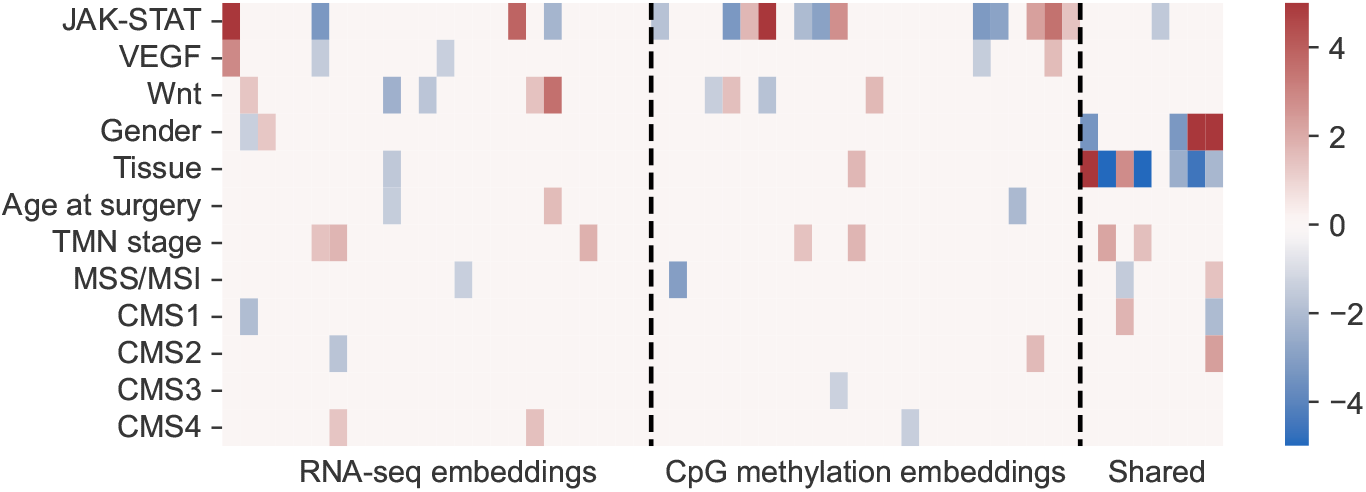
The heatmap shows the sign(coefficient) * -log10(P) values from each association test, where the coefficient gives the positive/negative direction. The columns are the 56 latent representation dimensions, and the rows are different annotations. The JAK-STAT, VEGF and Wnt represent gene set enrichment analysis of KEGG signaling pathways. The gender, tissue, age at surgery, TMN stage and MSS/MSI are clinical background information. The CMS1-4 labels were predicted from the RNA-seq data. Red color denotes positive correlation, and blue color denotes negative correlation.

#### 2.4.2 Interpretation of Omic-type Importances

In this section, we look deeper into the contributions of all four components of the latent representations on our performance metrics. In particular, we compute NMI and k-NN accuracy only using different omic-specific or shared representations, as well as their combinations. In other words, we drop the information relative to some omics, and observe the consequent drop (or increase) in performance. This allows us to understand whether the information relevant to the subtyping task is present in all of the omics, or concentrated within a specific subset. The result of those ablation studies on both MOCSS and our model is given in Fig. 7 and 12.

**Figure 7:**
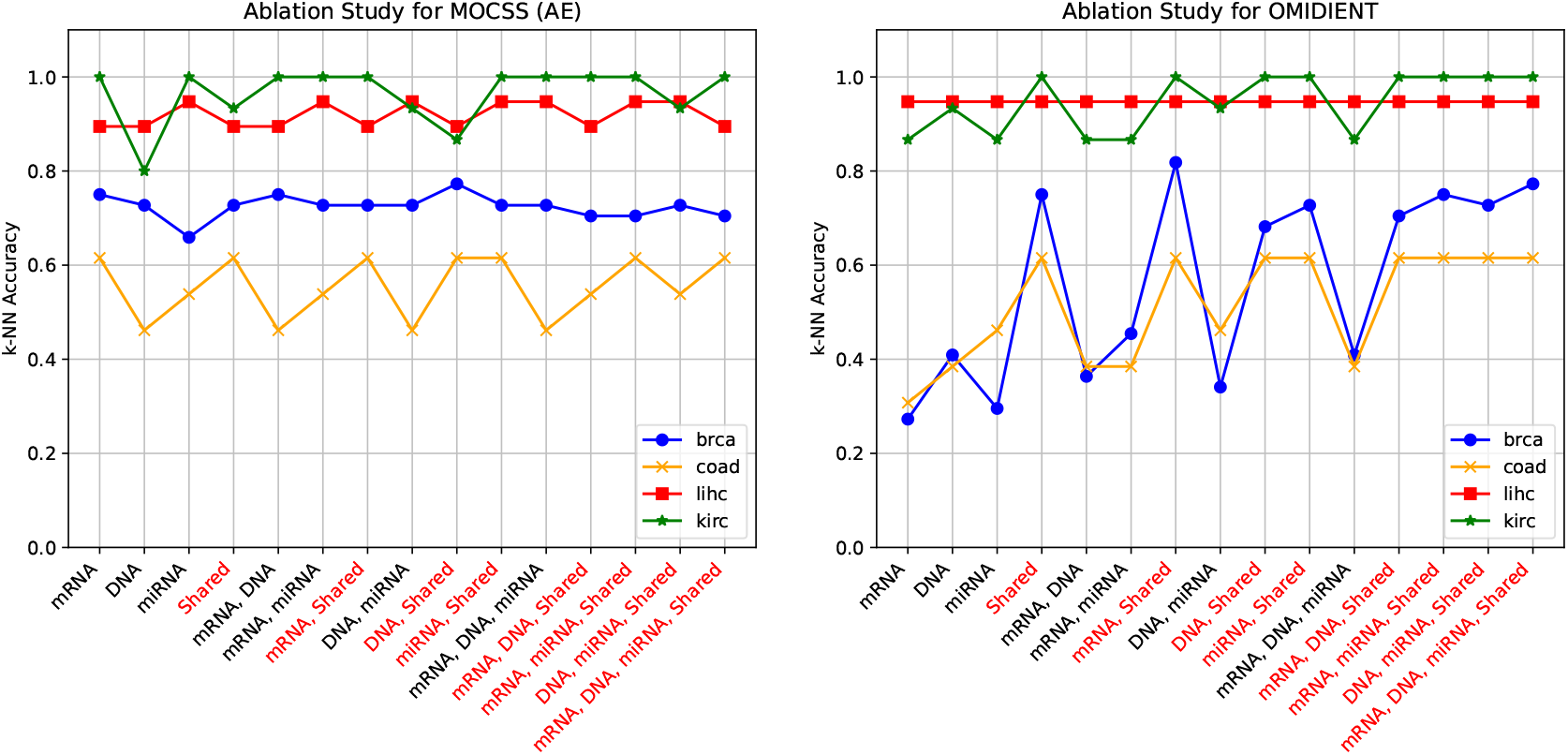
MOCSS’ and our method’s ablation study of different omic combinations based on k-NN accuracy. Highlighted values use the shared representation.

We observe that, for all four datasets, MOCSS (AE) reaches either its top or near-top performance with only a single omic available (mRNA for brca, coad and kirc, and miRNA for lihc). This suggests that the information necessary for subtyping was not effectively disentangled between the shared and omic-specific sub-representations, with each omic-specific representation largely retaining the information from its respective modality. Our model, on the other hand, shows a clear improvement of predictive performance when using the shared sub-representation. While the easier datasets (lihc and kirc) show a similar behavior to MOCSS, with a relatively high performance regardless of the subset of representations used, the two others exhibit their best (or close-to-best) performance either when all specific representations are present, or when the shared representation is used.

In order to formalize these observations, we display the average relative influence that each omic has on the performance. This value attribution is computed through the Shapley value Shapley (1953), a well-used method in explainable machine learning Lundberg and Lee (2017).

The Shapley values for all sub-representations are provided in Fig. 8. There, we confirm our previous observation, where, in MOCSS, the shared representation plays a preponderant role only on coad, yet only marginally so. On the other hand, in our approach, we see that the shared representation is significantly more influential than the three specific ones on 3 of the 4 datasets. We interpret this as showing that our shared representation is more expressive than MOCSS, achieving powerful extraction and aggregation of the information from each omic.

**Figure 8:**
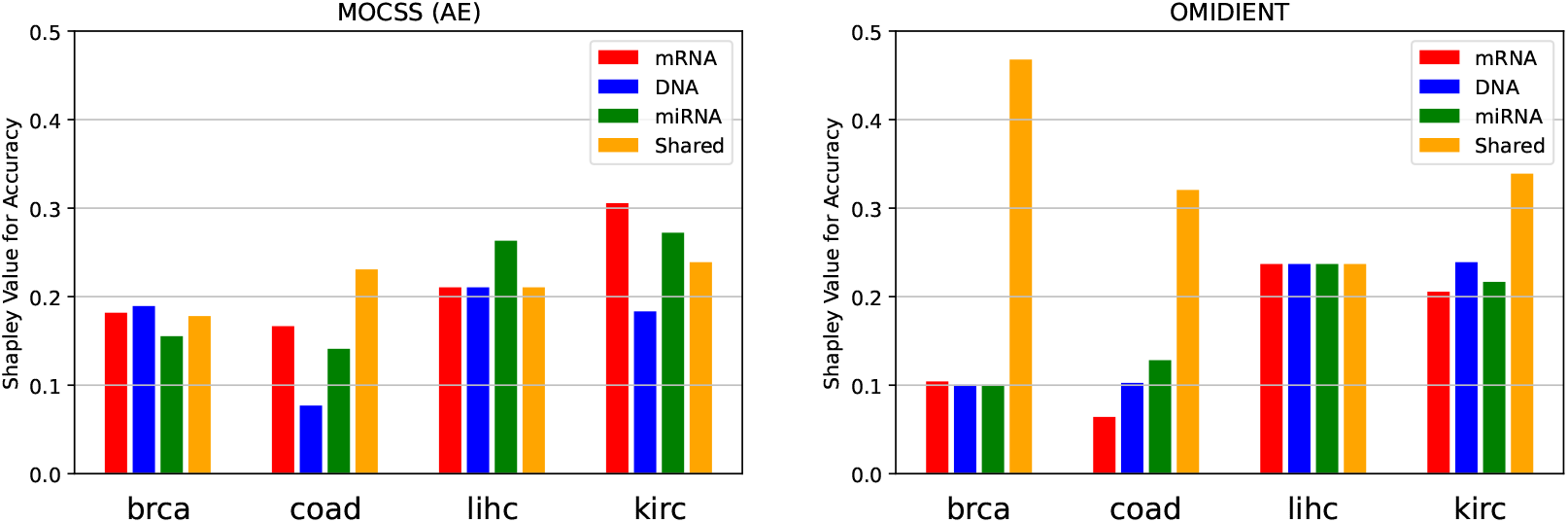
Shapley values for accuracy of k-NN

Finally, OMIDIENT, our proposed method, unlike MOCSS (AE), clearly uses different components of the whole representation for different purposes, i.e., classification and reconstruction. This is evidenced by our ablation study, which shows that OMIDIENT primarily relies on the shared representation for classification. The most informative signals across omics are concentrated in this shared part, consistent with the concept of shared latent spaces in multimodal integration You et al. (2022). Meanwhile, due to the architectural design of OMIDIENT, the reconstruction of each omic is handled independently through its omic-specific representation, ensuring that modality-specific details are preserved.

## 3 Methods

### 3.1 Framework

The deep learning architecture in OMIDIENT builds upon that of MOCSS Chen et al. (2023), with the shared goal of extracting both omic-specific and omic-shared information from multi-omics data. While MOCSS employs standard autoencoders to learn modality-specific features, OMIDIENT leverages a novel type of variational autoencoder framework, enabling it to model uncertainty and structure the latent space to better capture distinct tumor characteristics. We use a product of Dirichlet distribution to model the latent representations instead of the deterministic assignment of latent representations using a mere autoencoder; more details can be found in the next section, Probabilistic Graphical Model. For the omic-shared part, we use contrastive learning as in MOCSS; however, we do not use the loss function that MOCSS introduced. Namely, the orthogonal loss is not used in OMIDIENT to disentangle the omic-specific and omic-shared information. This choice is supported by our ablation study. As a result, we have three loss functions: reconstruction loss, contrastive loss and our novel loss for the product of Dirichlet variational autoencoder, which we explain in the next section.

### 3.2 Probabilistic Graphical Model

A standard variational autoencoder (VAE) Kingma and Welling (2014), shown in Fig. 9, learns a posterior distribution *q*_*ϕ*_(*Z*|*X*) (short form *q*(*Z*)) through a neural network, assuming that the prior distribution is *p*(*Z*) = *𝒩N* (0, *I*).

**Figure 9:**
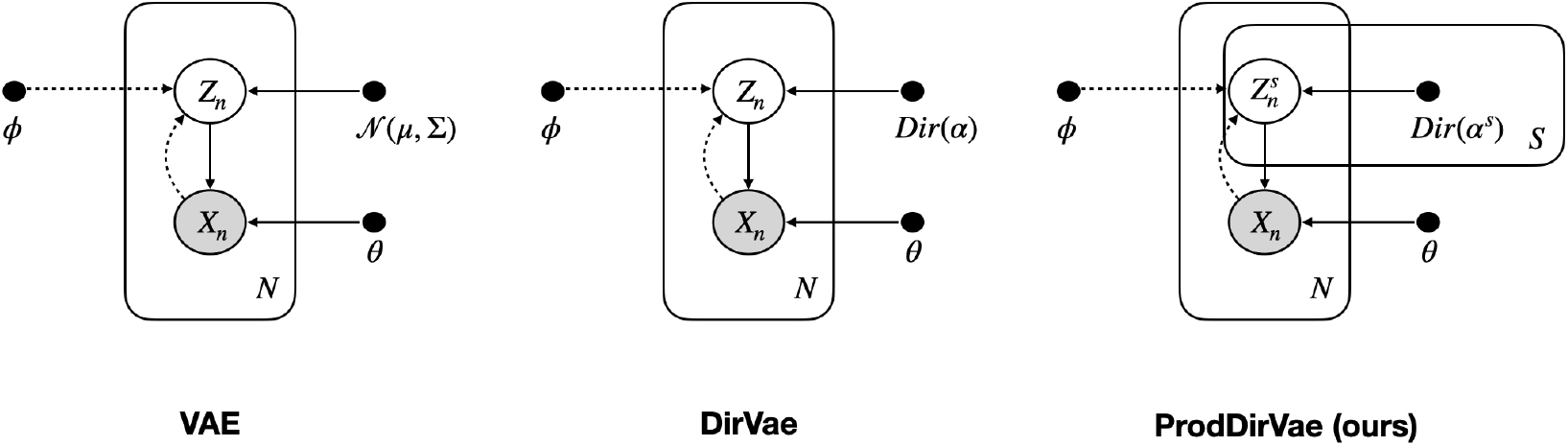
The graphical models of VAE, Dirichlet VAE (DirVae) and Product of Dirichlet VAE, ProdDirVae (our model) are shown.

Subsequently, other models have been proposed to model the prior as Dirichlet, Dirichlet Variational Autoencoders (DirVae), shown in Fig. 9, with various approximations; one method approximated Dirichlet by Laplace approximation Srivastava and Sutton (2017). Here we refer to it as LapDirVae, as shown in Eq. 1. A more recent method approximates Dirichlet samples by a Gamma distribution, instead of a Gaussian, by computing the inverse of the CDF of the Gamma distribution Joo et al. (2019):

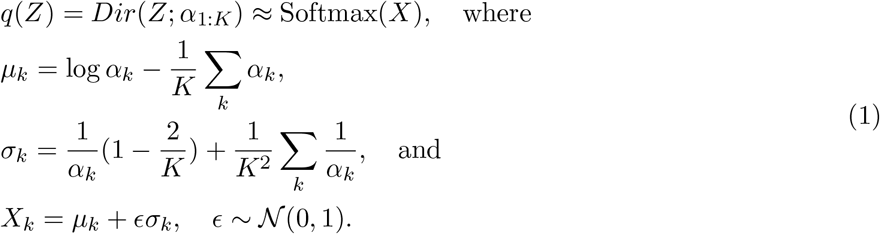

We propose *ProdDirVae*, shown in Fig. 9, a novel variational autoencoder that is governed by a product of *S* Dirichlet priors. Let *p*(*Z*) denote the prior, and let it follow *S* independent Dirichlet distributions, as shown in Eq. 2. Note that *S* is the number of groups into which the representations are decomposed. Moreover, *K* is the dimensionality of each group of representations, which represents the dimensionality of the Dirichlet distribution.

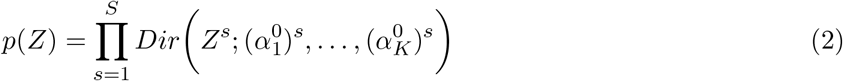

The motivation of this prior is three-fold: 1) the Dirichlet prior models the latent variable as a vector of categorical probabilities as opposed to the Gaussian distribution which assumes a smooth, unimodal distribution, not necessarily valid for tumors that may deviate nonsmoothly between individuals; 2) the product of Dirichlet decomposes the latent space into independent parts, i.e. enabling it to represent a variety of unrelated characteristics of a tumor; 3) Moreover, we fix the concentration parameters to be 0.5, which promotes sparsity of the tumor characteristics. This is the default Jeffreys prior for Dirichlet distribution with parameters 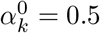.

We learn the posterior *q*(*Z*) by the encoder of the neural network. The posterior also takes the form of a product of Dirichlet distribution. For sampling the Dirichlet, we use the reparameterization trick by computing the inverse of the CDF of Gamma distribution as in Dirichlet VAE Joo et al. (2019), here we name it GamDirVae. Therefore, we have the following form shown in Eq. 3; for details of the approximation, see Appendix A.

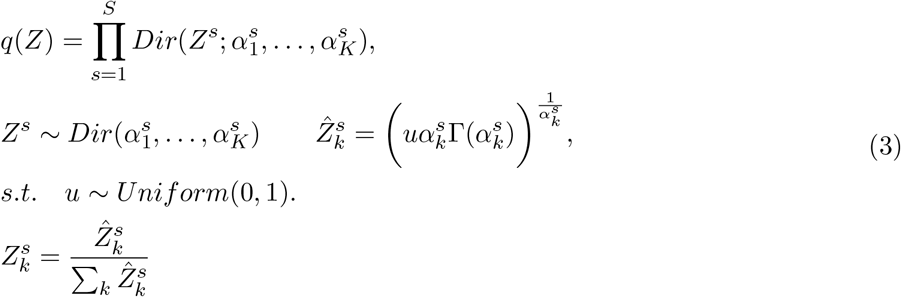

Given a prior Dirichlet distribution with parameters for each group 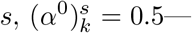 the Jeffreys prior— the Kullback–Leibler (KL) divergence between this prior and the approximate posterior with parameters 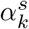 serves as the KL loss term in our model and is formulated as follows.

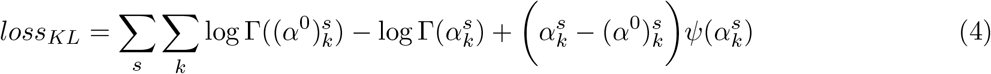

### 3.3 Encoder and Decoder

In this section, we present the novel components of our neural network architecture, i.e., the encoder and decoder for each omic modality; see Fig. 10 The encoder transforms the input omic data *X* into *S* groups of 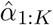. Subsequently, to ensure these parameters meet the positivity requirement of the Dirichlet distribution, i.e., positive concentration parameters, we apply a sigmoid activation function on them, resulting in concentration parameters *α*_1:*K*_. Using the concentration parameters and Gamma approximation trick, explained in Appendix A, the omic representation *Z* is computed. Our decoder begins by applying a softmax activation function to the temperature-scaled latent variable *Z*, i.e., *τ*^*−*1^*Z* with *τ* = ∑^*k*^ *Z*_*k*_. The scaling of the variable *Z* is intended to counteract the normalization effect inherent in the computation of the Dirichlet variable, allowing the model to instead take advantage of a sharper discrimination provided by the softmax function Zhang et al. (2023). The output of the softmax function serves as the basis for all the results reported in this paper. Ablation studies empirically demonstrate that this design choice leads to improved performance.

**Figure 10:**
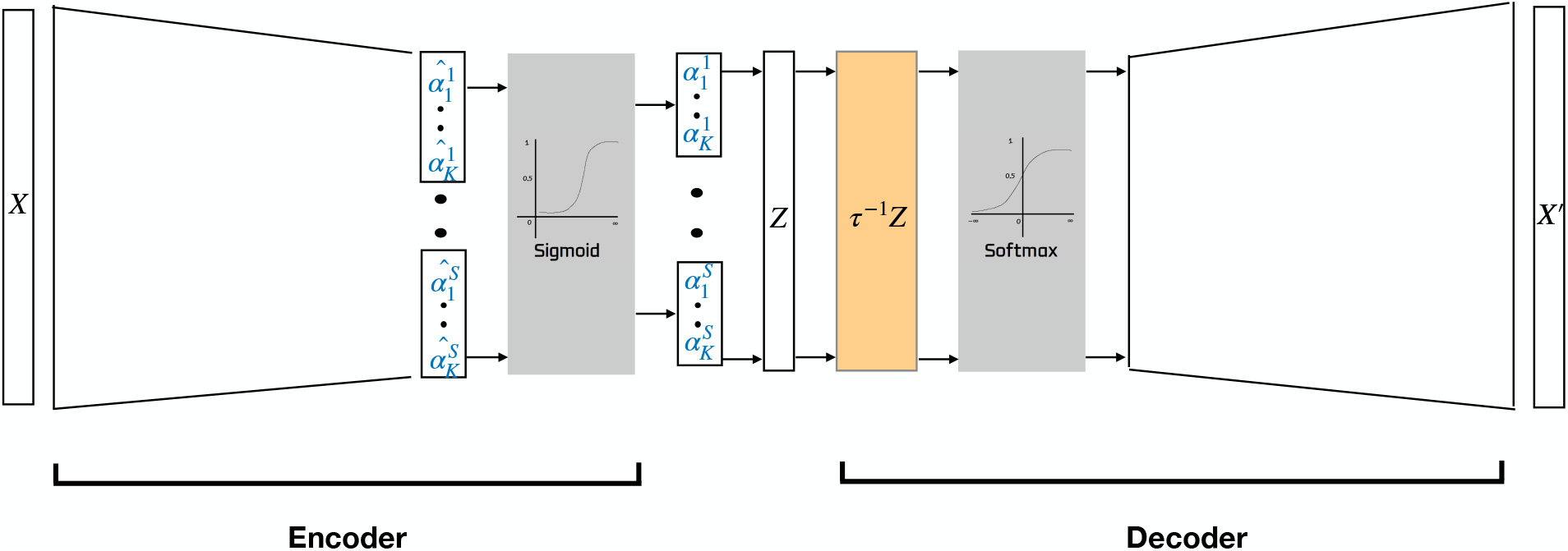
The overview of OMIDIENT’s encoder and decoder architecture.

### 3.4 Shapley Values

In order to formalize method interpretation, we display the average relative influence that each omic has on the performance. This value attribution is computed through the Shapley value Shapley (1953), a well-used method in explainable machine learning Lundberg and Lee (2017). Formally, assuming *n*_*o*_ omictypes, 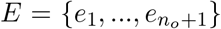 the set of sub-representations (one for each omic, and one shared). Then, we call *v* : 2^*E*^ *1→ ℝ*, the set function that returns the value of a metric (k-NN accuracy, K-Means NMI) when using only representations dimensions included in the set. In practice, the influence of sub-representation *e*_*i*_ on metric *v* is computed as:

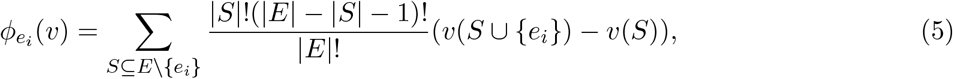

which is the average of the marginal contribution of *e*_*i*_ to each possible coalition of sub-representations. Since we have only four sub-representations (|*E*| = 4), it is feasible to use the Shapley value even if it has exponential complexity.

### 3.5 Setting

Table 4 summarizes the key hyperparameters used in our method (OMIDIENT) and the neural network based benchmarks. These include the shapes of the neural network layers, batch size, and the temperature parameter, which controls the model’s sensitivity to hard samples. It also lists the tuning hyperparameters: learning rate, weight decay, and the number of groups *S*, which determines the level of structured representation in our model.

**Table 4:** Hyperparameter Setting

| Hyperparameter | Setting |
| --- | --- |
| Shape of mRNA encoder-decoder layers | {512, 256, 128, 32, 32, 128, 256, 512} |
| Shape of DNA methylation encoder-decoder layers | {512, 256, 128, 32, 32, 128, 256, 512} |
| Shape of miRNA encoder-decoder layers | {256, 128, 64, 32, 32, 64, 128, 256} |
| Learning rate | [.0003, .0002, .0001, .0004, .0005, .0006] |
| Weight decay | [5e-4, 4e-4, 3e-4, 6e-4, 7e-4] |
| Number of groups for OMIDIANT | [2, 3, 4, 5] |
| Dimension of each group for OMIDIANT | 8 |
| Batch size | 32 |
| Temperature | 0.4 |

### 3.6 Dataset

We investigate the capacity of our method, OMIDIENT, with regard to representation learning by applying it to data from five different cancers: breast cancer (BRCA), colon adenocarcinoma cancer (COAD), colorectal cancer (CRC), liver cancer (LIHC), and kidney cancer KIR of type clear renal cell carcinoma (KIRC). The COAD, LIHC, and KIRC datasets are based upon data generated by The Cancer Genome Atlas (https://www.cancer.gov/tcga). The BRCA dataset obtained from MOCSS Chen et al. (2023), has labels indicating five subtypes within breast cancer, i.e., luminal A, luminal B, normal-like, basal-like, and HER2-enriched Parker (2009); Network (2012), each of which is classified by PAM50, a 50-gene signature Wang et al. (2021a). COAD dataset has labels indicating four subtypes within colon cancer, i.e., CIN, GS, microsatellite instability (MSI), and POLE Duan et al. (2021). Microsatellite instability (MSI) arises through defective DNA mismatch repair in the tumors, causing mutations particularly at repetitive DNA sequences called “microsatellites”. The LIHC dataset contains both cancer and normal samples Rappoport and Shamir (2018) and the KIRC dataset has two subgroups Xu et al. (2021). The CRC data was described in Ongen et al. (2014), and the published subset included the first 103 tumor-normal pairs, and the rest of the sample pairs were collected and processed in the same manner. CRC data has two types of annotations, one with four classes and another with two classes. MSI classification system is relevant for CRC prognosis and treatment Taieb et al. (2022). For the CRC with four classes, we used the following groups: 1) microsatellite instability (MSI), 2) microsatellite stability (MSS), 3) ambiguous tumors, and 4) normal samples. For the CRC with consensus molecular subtypes (CMS) grouping Guinney et al. (2015), we used the following groups: 1) CM1 group containing microsatellite instability immune, 14%, hyper-mutated, microsatellite unstable, and strong immune activation, and 2) other CMS groups. The set of CRC samples was described in Ongen et al. (2014). The published subset included the first 103 tumor-normal pairs, and the rest of the sample pairs were collected and processed in the same manner. For all datasets except CRC, three types of omics data, i.e., mRNA expression data (mRNA), DNA methylation data (meth), and miRNA expression data (miRNA), are used to provide integrative and complementary information on each cancer dataset. For CRC, mRNA and DNA methylation are available, and the top 10000 features with the largest variance in each omic are chosen. The BRCA dataset is preprocessed with noise reduction Wang et al. (2021a), which results in a more refined list of omic features. Feature engineering is not applied to the rest of the datasets. The information on the number of features in each omic data for each cancer and the number of cancer samples is listed in Table 5.

**Table 5:** Summary of datasets with the number of tumor samples and ground-truth classes

| Dataset | mRNA expression | miRNA expression | DNA methylation | Samples | Classes |
| --- | --- | --- | --- | --- | --- |
| BRCA | 1000 | 503 | 1000 | 875 | 5 |
| COAD | 17260 | 17261 | 19052 | 260 | 4 |
| CRC | 10000 | N/A | 10000 | 309 | 4, 2 |
| LIHC | 20530 | 1046 | 5000 | 378 | 2 |
| KIRC | 58315 | 1879 | 22928 | 289 | 2 |

## 4 Discussion

The advancement of omics technologies has improved personalized medicine, particularly transforming cancer research and treatment through more precise diagnosis, subtyping, and therapy selection. Despite the availability of more data annotations than before, multi-omics integration methods that require these labels (supervised approaches) lose the ability to discover new phenotypes or traits (e.g., disease diagnosis, grading of tumors, and cancer subtypes) since they become trained for certain pattern recognition introduced by the labels. To this end, we proposed OMIDIENT, an unsupervised (without using labels) multi-omics integration framework that is capable of finding novel molecular or clinical subtypes. OMI-DIENT is based on deep generative models, where we learn comprehensive embeddings representing both omic-specific and omic-shared information of multi-omics data. This compressed and informative representation can subsequently be used for various downstream tasks such as cancer subtyping. OMIDIENT also demonstrates strong capability in reconstructing missing values across omics datasets. To sum-marize, OMIDIENT is a novel multi-omics representation learning framework based on deep generative models with superior performance in typical use-cases compared to other state-of-the-art unsupervised approaches.

Unlike similar generative models, OMIDIENT can capture structured biological or functional relationships by dividing omic features into distinct groups. The use of groups enables the separation of embeddings, yielding a more structured and modular representation of multi-omics data that captures its inherent complexity. While we only demonstrated the capability of OMIDIENT with respect to subtyping, classification, data reconstruction, and limited biological interpretability, OMIDIENT allows for improving the expressiveness and potential interpretability of the learned representations due to its novel notion of feature grouping. Further investigation can be conducted to identify explicit groups within the learned embeddings to infer gene sets or any phenotypic categories. As such, the contribution of each biological module can be analyzed both independently and in combination.

Finally, despite the fact that we utilized mRNA expression, DNA methylation, and miRNA expression data for the multi-omics integration in this paper, the framework can be easily extended to accommodate additional data types.

## Author contributions statement

N.S., P.M., and L.A. designed the study. N.S. and P.M. conceived the model, framework, and evaluation tools. N.S., N.V., and L.A. collected and assembled the data. N.S. implemented all the modules for the novel method. N.S. and R.B. provided the method interpretation section. N.S., N.V. and K.R. performed biological interpretations. N.S., A.G., and N.V. performed benchmarking evaluations. All authors contributed to the final writing and approval of the manuscript.

## A Dirichlet Representations

In this section, we review the details for the reparameterization trick in modeling the latent representation with a Dirichlet distribution Joo et al. (2019); this is also useful for our novel model: the product of Dirichlet distributions. Firstly, we know that a Dirichlet distributed values with *K* dimensions can be formulated as *K* normalized Gamma distributions, i.e., *Z*_*k*_ *∼Gam*(*α*_*k*_, 1) and *normalized*(*Z*_1_, …, *Z*_*K*_) *∼Dir*(*α*_1_, …, *α*_*K*_). Ffor sampling Gamma values, we use the Inverse CDF Method, or Probability Integral Transform, which states that for any random variable *Z* with its CDF, *F* (*Z*), we can sample *Z* by first sampling *u ∼Unif* (0, 1), and then computing *Z* = *F*^*−*1^(*u*) Bishop (2006). Given 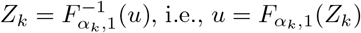, we write the approximated Gamma CDF for small values of *Z*_*k*_—since these will be normalized for the Dirichlet samples we are not constrained to small values in the final step.

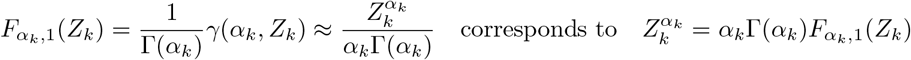

Computing the inverse of the above, we achieve

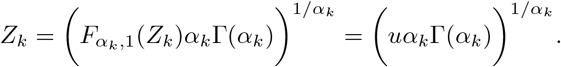

Letting *C* denote *uα*Γ(*α*), Figure 11 illustrates the final reparameterization in the Dirichlet networks. Note that we use sigmoid to ensure *α* vector of concentration parameters stays positive, and we normalize the sampled Gamma values so that the samples have the Dirichlet distribution.

**Figure 11:**
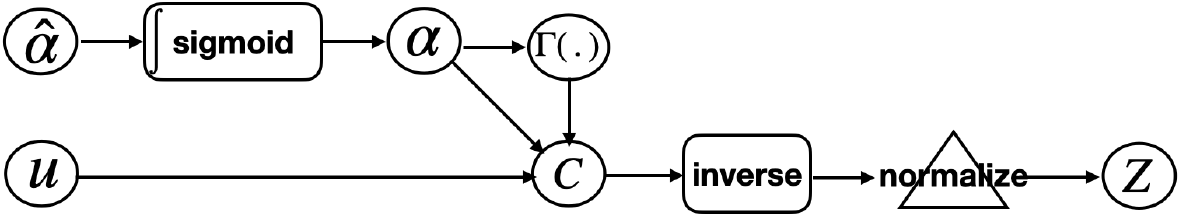
The Dirichlet Reparameterization of variable *Z*.

## B Model Explainability

### B.1 Ablation Studies

Fig. 12 depicts the result of the ablation studies for our model, OMIDIENT. Fig. 13 to 17 depicts the result of the ablation studies for other models and all diseases.

**Figure 12:**
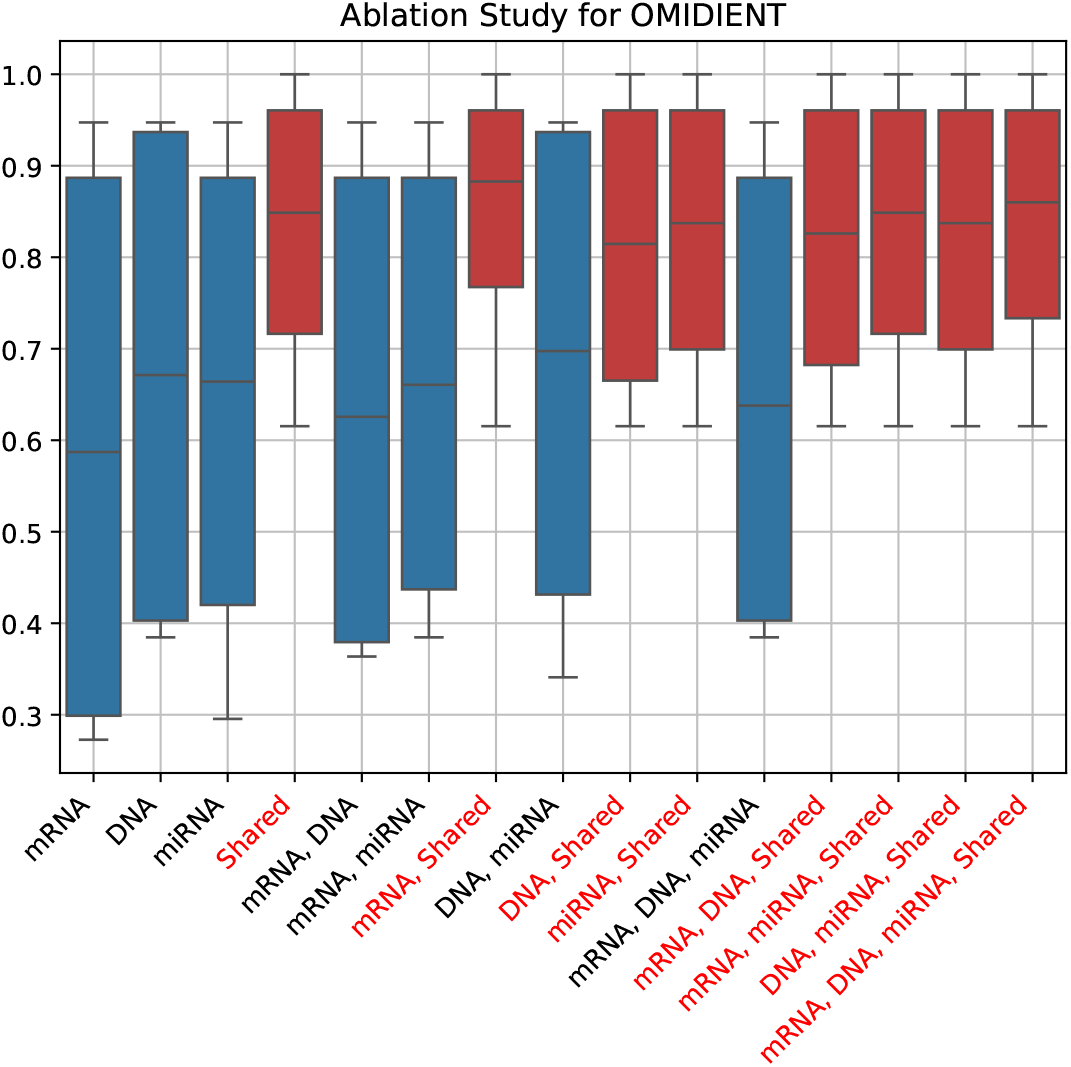
Boxplot view of figure 7. The red boxes use the shared representation.

**Figure 13:**
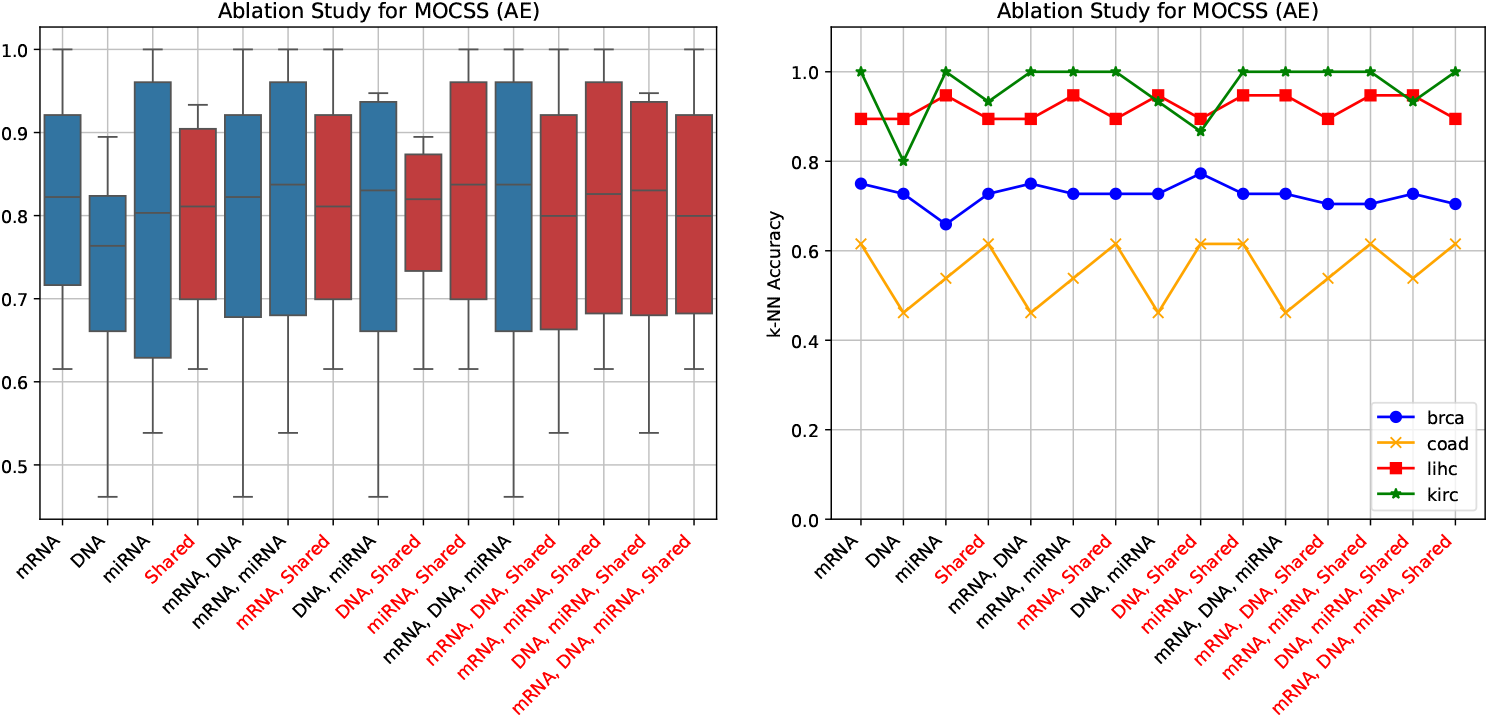
Ablation results values AE (MOCSS)

**Figure 14:**
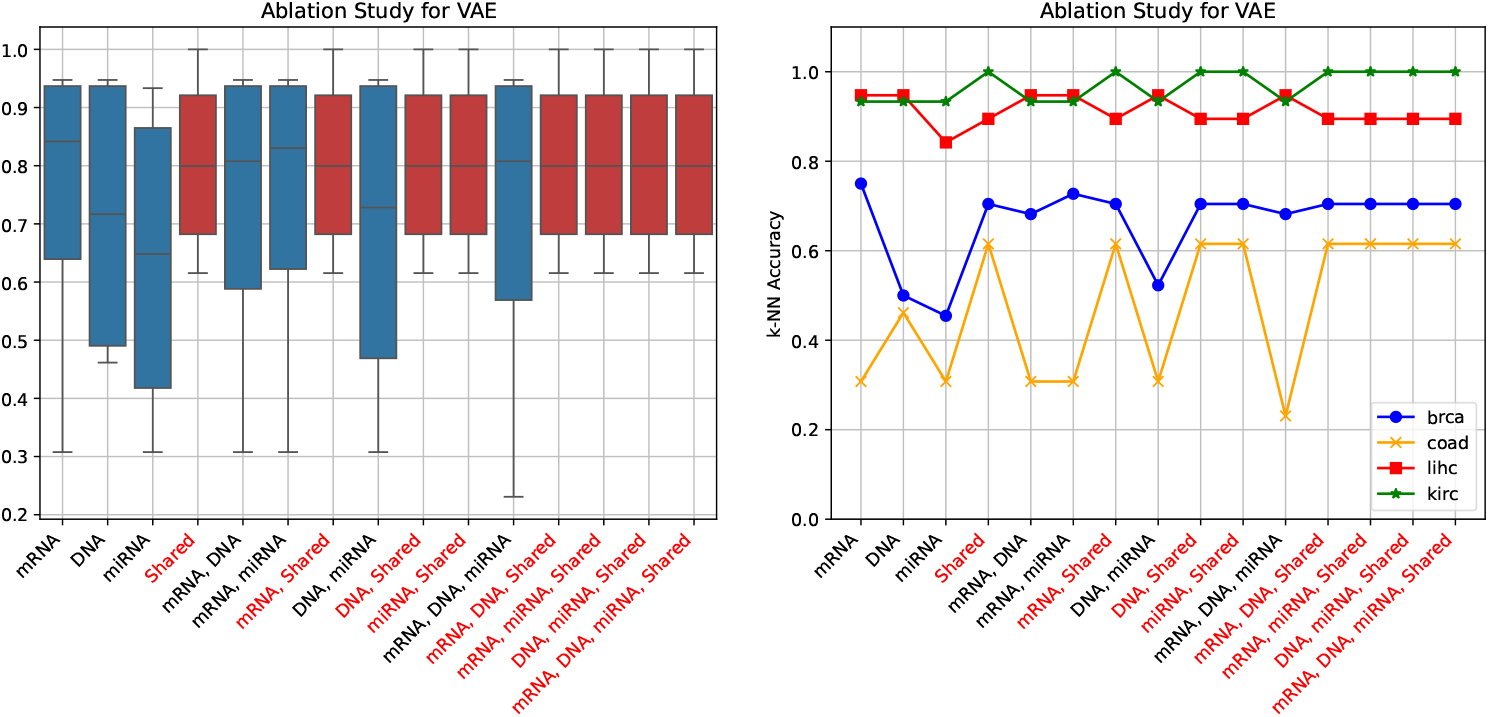
Ablation results values for accuracy of VAE

**Figure 15:**
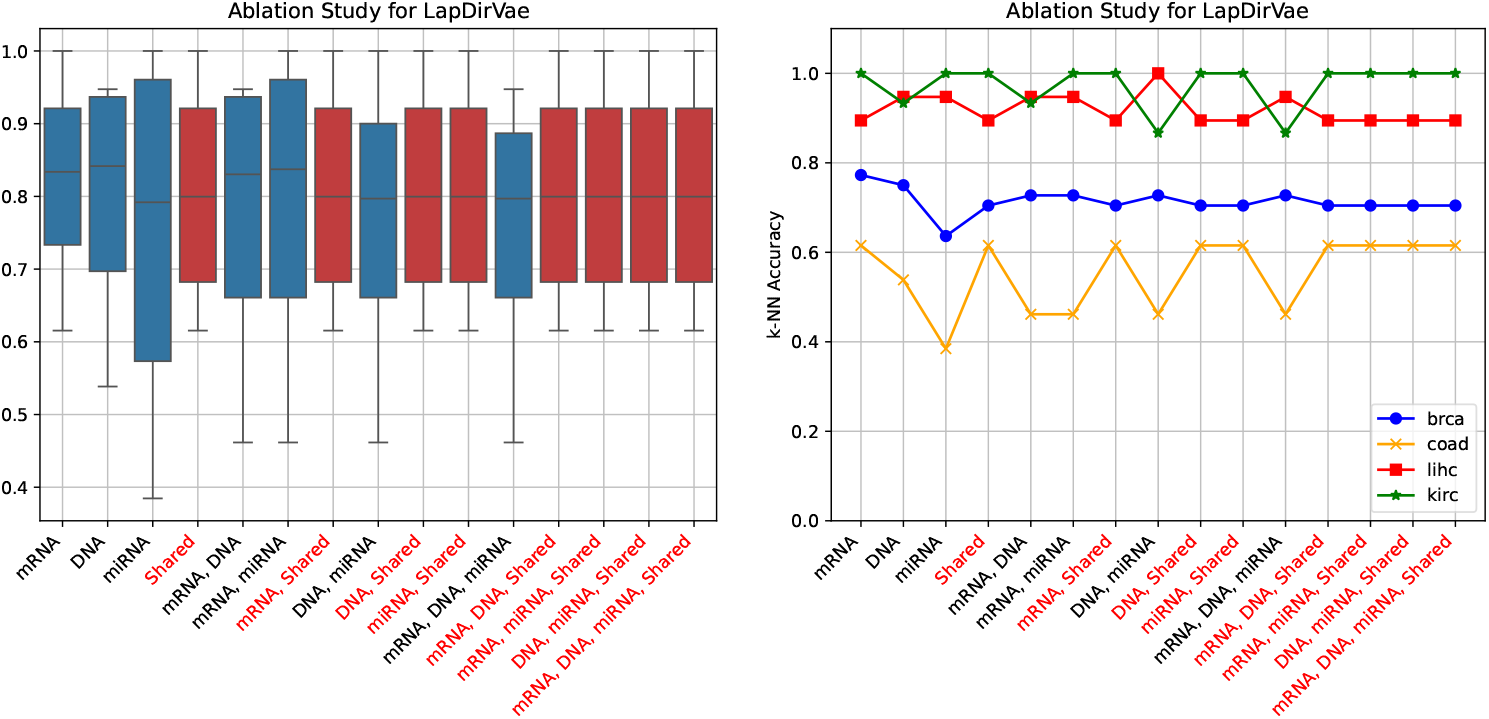
Ablation results values for accuracy of LapDirVAE

**Figure 16:**
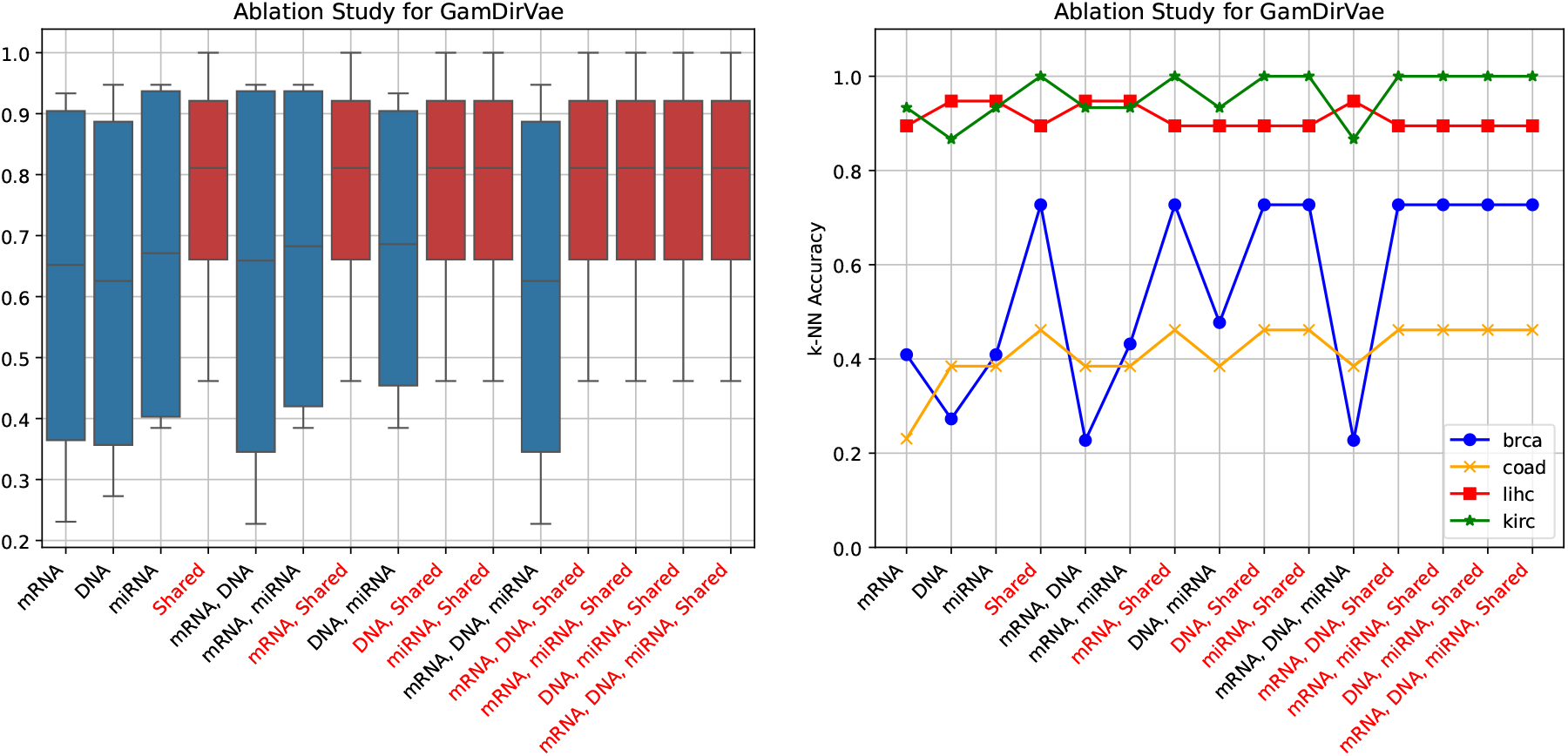
Ablation results values for accuracy of GamDirVae

**Figure 17:**
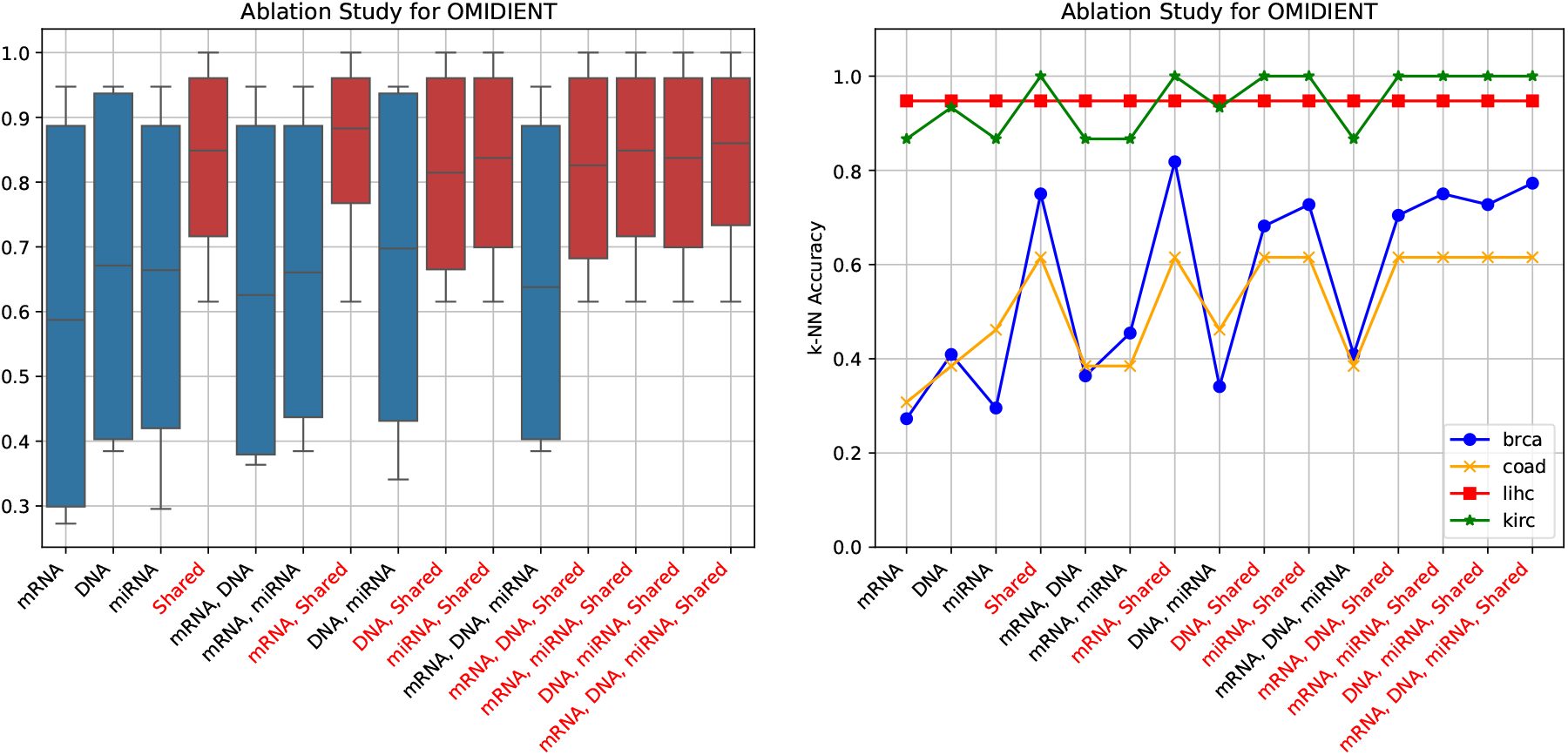
Shapley values for accuracy of OMIDIENT

### B.2 Shapley values

Fig. 18 depicts the Shapley values of the different sub-representations for all models and all diseases.

**Figure 18:**
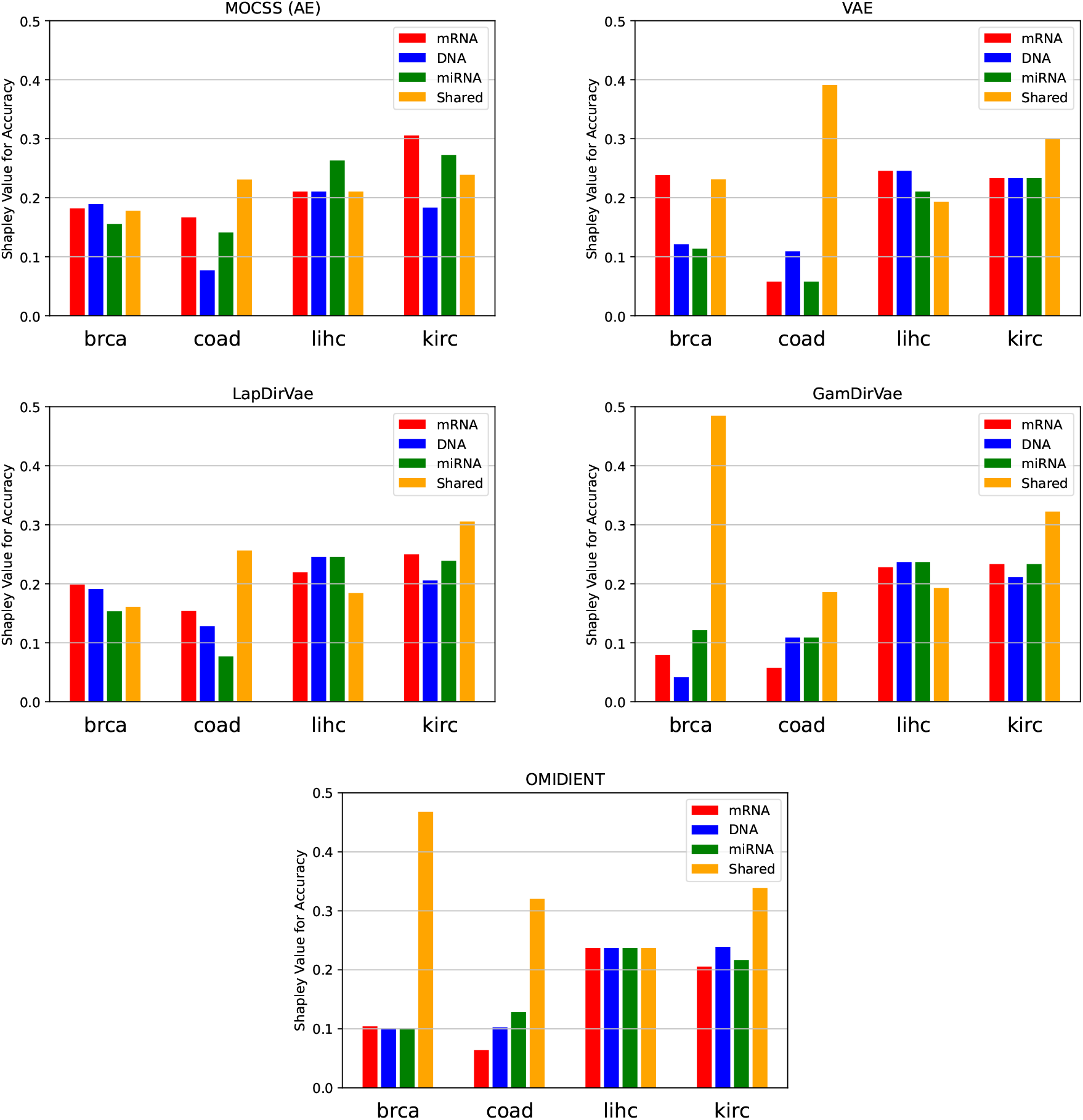
Shapley values for accuracy of k-NN

